# Nonlinear System Identification of Neural Systems from Neurophysiological Signals

**DOI:** 10.1101/2020.08.09.243253

**Authors:** Fei He, Yuan Yang

## Abstract

The human nervous system is one of the most complicated systems in nature. Complex nonlinear behaviours have been shown from the single neuron level to the system level. For decades, linear connectivity analysis methods, such as correlation, coherence and Granger causality, have been extensively used to assess the neural connectivities and input-output interconnections in neural systems. Recent studies indicate that these linear methods can only capture a small amount of neural activities and functional relationships, and therefore cannot describe neural behaviours in a precise or complete way. In this review, we highlight recent advances in nonlinear system identification of neural systems, corresponding time and frequency domain analysis, and novel neural connectivity measures based on nonlinear system identification techniques. We argue that nonlinear modelling and analysis are necessary to study neuronal processing and signal transfer in neural systems quantitatively. These approaches can hopefully provide new insights to advance our understanding of neurophysiological mechanisms underlying neural functions. These nonlinear approaches also have the potential to produce sensitive biomarkers to facilitate the development of precision diagnostic tools for evaluating neurological disorders and the effects of targeted intervention.

## I. INTRODUCTION

The human nervous system is a complicated network comprised of more than 10 billion neurons, with trillions of synapses connecting them. Neuronal information processing is complex at different levels, from the microscopic pre- and post-synaptic cellular interactions to the macroscopic interactions between large populations of neurons in, for instance, sensory processing and motor response [1]. The behaviour of a single neuron is highly nonlinear, showing a step-like ‘none-or-all’ firing response [2], while the behaviour of neurons in a population could be relatively similar. Therefore, the nonlinear response of each individual neuron may be smoothed out by the distribution of membrane thresholds across the population, known as the pool effect [3]. This effect typically occurs in a mono-synaptic neural system such as the corticospinal tract where the supraspinal motor command is linearly transferred to the motor output due to the pool effect of motor units [4]. However, multi-synaptic neural systems, such as the somatosensory system, have been reported highly nonlinear, showing harmonic responses to periodic stimuli [5–8]. Cross-frequency coupling in the corticothalamic interactions has also been reported when characterising essential tremor [9]. Nonlinear behaviours in neural systems are thought to be associated with various neural functions, including neuronal encoding, neural processing of synaptic inputs, communication between different neuronal populations and functional integration [10–13].

Various functional and effective connectivity measures have been developed [14] to characterise such linear and nonlinear functional integration in neural networks, from large-scale neurophysiological signals. These signals, such as electroencephalogram (EEG) and magnetoencephalogram (MEG) from the brain, electromyogram (EMG) from muscles, measure neural activities from macro-scale neuronal populations. While functional connectivity measures, e.g. correlation, coherence, mutual information, only quantify the undirected statistical dependencies among signals from different areas, effective connectivity attempts to quantify the directed causal influences of one neural system over another, either at a synaptic or population level [15]. Mostly, effective connectivity measures are based on models of neural interactions or coupling (although there exist model-free measures like transfer entropy [16]) and is often time-dependent (dynamic). Therefore, effective connectivity has a strong link with dynamic modelling, also known as system identification in control systems theory [17–19], and corresponding model-based causality analysis.

Nonlinear system identification techniques have been formally applied to study neuronal information processing and neural systems since the 1970s. Some pioneering work includes: the nonlinear dynamic modelling of the retinal neuron chains in receptive-field responses [20, 21], the identification of nonlinear synaptic interactions [22], the identification of neural systems using stimulus-response and white-noise approach [23–25], the development of nonlinear systems analytic approach based on functional power (or Volterra/Wiener) series to study central nervous system function and hippocampal formation [26, 27], and nonlinear identification of stretch reflex dynamics [28]. Until now, many linear and nonlinear system identification methods have then been proposed and developed in the neuroscience context. Nevertheless, recent studies indicate that linear methods can only capture a small amount of neural activities and functional relationships, and therefore cannot describe neural behaviours in a precise or complete way [29, 30]. Nonlinear approaches provide us with useful tools to explore the nonlinear nature of neural systems [10]. In this review article, we highlight the need and recent advances in nonlinear system identification of neural systems, as well as novel neural connectivity analysis methods based on nonlinear system identification techniques. A diagram that summarises the linear and nonlinear functional and effective connectivity measures and their links with system identification is provided in Figure 1.

**FIG. 1:**
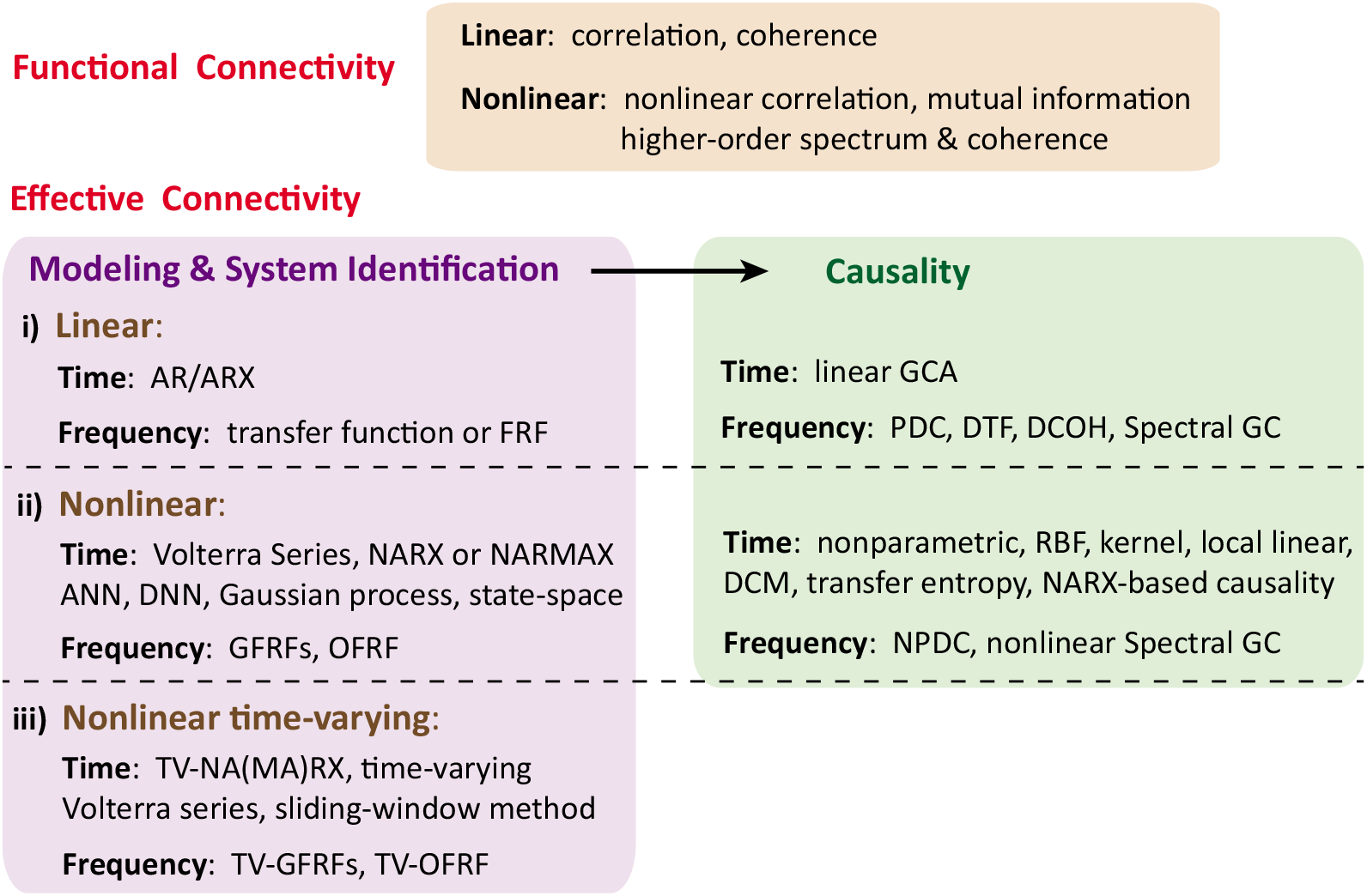
Overview of the linear and nonlinear functional and effective connectivity (causality) measures and their links with system identification methods. The linear functional connectivity, linear system identification and linear causality measures are first reviewed in Section III. The nonlinear and nonlinear time-varying system identification approaches (in both time and frequency domains) are then investigated in Section IV. The recently proposed nonlinear function connectivity measures and nonlinear causality measures (based on nonlinear system identification) are introduced in Section V. The abbreviations in the diagram are defined as: autoregressive (AR), autoregressive with exogenous input (ARX), frequency response function (FRF), nonlinear autoregressive (moving average) model with exogenous inputs (NAR(MA)X), artificial neural networks (ANN), deep neural networks (DNN), generalised frequency response function (GFRF), output frequency response function (OFRF), time-varying (TV), Granger causality analysis (GCA), partial directed coherence (PDC), directed transfer function (DTF), spectral Granger causality and directed coherence (DCOH), Granger causality (GC), dynamic causal modelling (DCM), nonlinear partial directed coherence (NPDC).

## II. NONLINEARITY IN THE NEURONAL LEVEL AND NEURAL SYSTEMS

At a single neuron level, the action potential spike is the principal basis of information encoding, which allows signal transmission across different neuronal populations [13]. The spike timing is thought to be associated with the coding scheme in neural systems [31]. The nonlinear nature of the neuronal process of synaptic input influences the temporal firing behaviour of individual neurons. Different types of neurons have their own repertoire of ion channels that are responsible for their characteristic nonlinear firing patterns and associated neural functions. For example, persistent inward currents mediated by their voltage-gated sodium and calcium channels are an important source of the nonlinear behaviour of spinal motoneurons. They are instrumental in generating the sustained force outputs required for postural control [32]. Activation of the L-type calcium channels in nigral dopaminergic neurons results in intrinsic bursting behaviour [33], exhibiting low-dimensional determinism and likely encodes meaningful information in the awake state of the brain [34]. The nonlinearity of the neuronal transfer function mediated by its component ion channels can generate various types of nonlinear output patterns such as harmonic, subharmonic and/or intermodulation of input patterns.

Despite plenty of knowledge of the nonlinear behaviour of a single neuron, the input-output relation at the neural system level is yet to be understood entirely. The system-level neural response is a composite output of collective neuronal activities from a large number of neurons. In a neuronal population, the pool effect can reduce the nonlinearity generated from each individual neuron, by smoothing the neuronal dynamics from a scale of milliseconds (spikes) to 10 milliseconds (local field potentials) or to 100 milliseconds (large-scale neurophysiological activities or signals such as EEG) [35]. Such effects have been previously demonstrated through both a computational model and an in vivo study in the human motor system, where the motor command can be transmitted linearly via the mono-synaptic corticospinal tract when more than five motoneurons are activated [4]. However, a small amount of nonlinearity may still be present [36]. A recent study simulated nonlinear neuronal dynamics on a large-scale neural network that captured the inter-regional connections of neocortex in the macaque. The authors applied information-theoretic measures to identify functional networks and characterized structure-function relations at multiple temporal scales [37]. The nonlinearity in each neuronal population can cumulatively increase if the system involves multiple synaptic connections [38]. A recent study in hemiparetic stroke shows that the nonlinearity in the motor system increases due to an increased usage of multi-synaptic indirect motor pathways, e.g. cortico-reticulospinal tract [39], following damage to the mono-synaptic corticospinal tract [40].

Assessing the input-output relation in neural systems, e.g. sensory, motor and cognitive processes, is essential to a better understanding of the nervous system. For instance, it could help to gain a better insight into the normal and pathological neural functions. It is well known that a linear system generates only iso-frequency interactions between an input and the output, e.g. the coupling of neuronal oscillations at a specific frequency band [41]. For decades, correlation and coherence measures have been used to identify the linear interaction in neural systems. More recently, various studies indicate the input-output neural interactions can cross different frequency components or bands, which is named cross-frequency coupling [42–45] and is a distinctive feature of a nonlinear system. In the following sections, we review both the linear and nonlinear approaches for identifying neural systems and associating neural connectivity, especially from a system identification perspective.

## III. LINEAR CONNECTIVITY AND SYSTEM IDENTIFICATION

The nervous system is a highly cooperative network composed of different groups of neurons. Neural connectivity, i.e., the synchronization of neural activity across these groups, is crucial to the coordination among distant, but functionally related, neuronal groups [46]. Linear neural connectivity can be assessed by determining the signal correlation or causality between the recorded neural signals. This section reviews commonly used linear connectivity, system identification methods and their interconnections in studying neural systems.

### A. Correlation and coherence

The most widely used measure of interdependence between two time series in the time domain is the crosscorrelation function [14], which measures the linear correlation between two signals or stochastic processes *X* and *Y* with discrete observations *x*(*t*) and *y*(*t*), at *t* = 1, 2,…, *N*, as a function of their delay time:

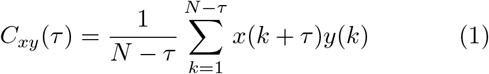

where *N* is the number of samples and *τ* the time lag between two signals. This function ranges from −1 (complete linear inverse correlation) to 1 (complete linear direct correlation). The value of *τ* that maximizes this function can be used to estimate the linearly related delay between signals. The well-known Pearson correlation coefficient is equal to *C_xy_*(*τ*) when *τ* = 0. The linear dependence between two signals in the frequency domain is usually measured by the spectral coherence. The coherence between two signals at frequency *f* is defined as:

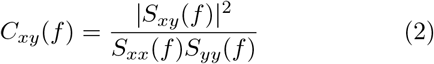

where *S_xy_*(*f*) is the cross-spectral density between *x* and *y*, and *S_xx_*(*f*) and *S_yy_*(*f*) the auto-spectral density of x and y respectively. The cross-spectral and auto-spectral densities are the Fourier transforms of the crosscorrelation and auto-correlation functions of the two signals. Values of coherence are always between 0 and 1.

The correlation and coherence measures have been widely applied to EEG, MEG or EMG signals to characterise the neuronal interactions, from the firing of cortical neuron spike trains to complicated neural systems (for reviews see [14, 47]).

### B. System identification and causality

Unlike functional connectivity, effective connectivity emphasises on the directional causal influences between neural areas or signals. Here, we first introduce the classical Granger causality and its link with the time-domain linear system identification, i.e. regression models of bivariate time series. The frequency-domain causality measures can then be linked with the frequency response function of linear systems.

Considering two signals or variables *X* and *Y*, the interactions of the signals can be described by bivariate linear autoregressive with exogenous input (ARX) models jointly,

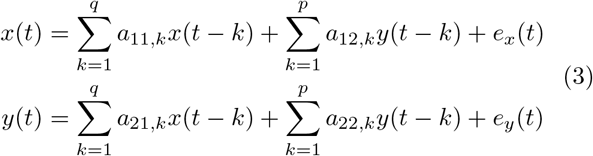

where *p* and *q* are the model order of *y* and *x* regressors; *e_x_*(*t*) and *e_y_*(*t*) are the uncorrelated model prediction errors over time. A linear causal influence from *X* to *Y* defined by Granger can be expressed as a log ratio of the prediction error variances of the corresponding restricted (AR) and unrestricted (ARX) models:

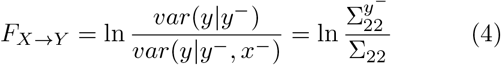

where *x*^−^ and *y*^−^ denotes contributions from lagged input and output terms, respectively; *y*^−^_22_ denotes the variance of *e_y_* when there are only regression terms of *Y*. The linear ARX models (3) can be re-written in matrix form and mapped to the frequency domain by Fourier transformation:

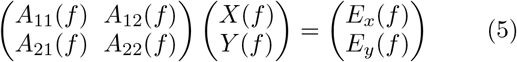

where the components of the coefficient matrix **A**(*f*) are 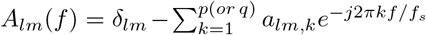 with *f_s_* the sampling frequency and *δ_lm_* the Kronecker delta function. We can re-write the above equation by inverting the coefficient matrix **G**(*f*) = **A**^−1^(*f*) and moving the so-called transfer function matrix **G**(*f*) to the right-hand-side the equation. Different frequency-domain Granger causality measures, such as partial directed coherence (PDC), directed transfer function (DTF), spectral Granger causality and directed coherence (DCOH) [48–50], can then be expressed as a function of the elements of either the coefficient matrix **A**(*f*) or the transfer function matrix **G**(*f*) (Baccala and Sameshima, 2001; Chicharro, 2011). By dividing both sides of (5) with the corresponding diagonal elements in the coefficient matrix **A**, the off-diagonal elements in the transformed coefficient matrix are actually related to the negative frequency response functions (FRFs) of linear ARX systems, if one signal is treated as the input while the other is treated as the output. For instance,

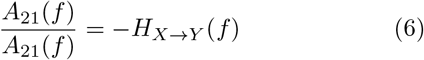

the FRF, *H*_*X*→*Y*_(*f*), describes the input-output relationship, i.e., with input *X* and output *Y*, of the (noise-free) ‘system’ in the frequency domain. It is also known as the ‘transfer function’ in linear system theory. Frequency-domain Granger causality measure, e.g. PDC, can be expressed as a function of the FRFs of the corresponding linear ARX and AR models:

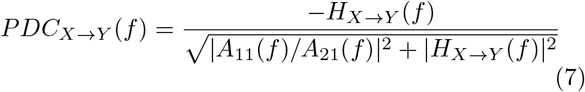

Establishing such a link between the causality measures and linear system identification, in both time and frequency domains, is crucial to the further development of nonlinear model-based causality measures via nonlinear system identification, which will be investigated in Section IV and V.

### C. Limitation of linear approaches on identifying neural system

Linear connectivity and system identification allow the assessment of communication between neuronal populations at the same oscillatory frequency band or similar neuronal firing patterns. The applications of linear approaches have been thoroughly reviewed previously [51, 52]. However, it is not clear how much information is missing when using the linear approach since the behaviour of various neural systems can be highly nonlinear [10, 11]. When one uses a linear measure to investigate a neural system, the nonlinear neural interaction is ignored, especially between the neuronal populations which have very different mean firing rates such as the central nervous system and the periphery. A recent study reported that in the human somatosensory system over 80% of the cortical response to wrist joint sensory input comes from nonlinear interactions, where a linear model explains only 10% of the cortical response [29]. Therefore, nonlinear connectivity and modelling approaches are needed to investigate neural systems in a complete way.

## IV. NONLINEAR SYSTEM IDENTIFICATION OF NEURAL SYSTEMS

It is often impossible to derive a mechanistic model of a neural system, due to the complexity of the underlying biological process and many unobservable state variables. In this section, we focus on the generic nonlinear model representations of a single-input and single-output (SISO) neural dynamic system, its identification process in the time domain, and corresponding frequency-domain analysis. We first investigate the identification of nonlinear time-invariant systems, and then time-varying nonlinear systems.

### A. Time-domain nonlinear system identification

#### 1. Volterra series

The Volterra series model is a direct generalisation of the linear convolution integral and provides an intuitive representation for a nonlinear input-output system. The output *y*(*t*) of a SISO nonlinear system can be expressed as a Volterra functional of the input signal *u*(*t*):

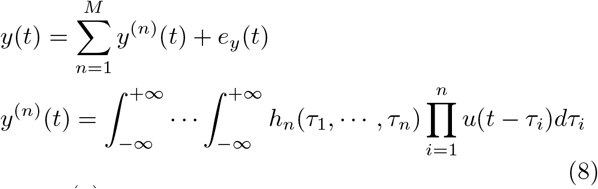

where *y*^(*n*)^(*t*) is the *n*th-order output and *M* is the maximum order of the system’s nonlinearity; *h_n_*(*τ*_1_, ⋯, *τ_n_*) is the *n*th-order impulse response function or Volterra kernel, which describes nonlinear interactions among *n* copies of input and generalises the linear convolution integral to deal with nonlinear systems. Neurobiologically, Volterra series can be directly interpreted as the effective connectivity - ‘the influence that one neural system exerts over another, either at a synaptic or population level’ [53]. The first-order kernel describes the linear ‘driving’ efficacy or linear synchronous interactions, and the second- or higher-order kernels represent the ‘modulatory’ influence or asynchronous interactions [10]. The Fourier transform of the first-order kernel is the FRF (or transfer function) and describe the interactions in the same frequencies, while the frequency-domain counterparts of the higher-order kernels are the GFRFs (to be discussed in Section IV B) which quantify the cross-frequency interactions.

Practically, to deal with a large number of Volterra series coefficients, a regularization strategy is often employed in the estimation procedure [54]. Volterra model has been widely used in physiological systems, including neural systems, modelling. Some recent examples include the study of nonlinear interactions in the hippocampal-cortical neurons [55], in the spectrotemporal receptive fields of the primary auditory cortex [56], in the sensory mechanoreceptor system [29], in the human somatosensory system (i.e. the cortical response to the wrist joint sensory input) indicating the dominance of nonlinear response [30], in multiple-input and multiple-output (MIMO) spiking neural circuits [57] and hippocampal memory prostheses [58]. The Volterra model also has a strong theoretical link with the NARMAX model [59] and the dynamic causal modelling [60] to be discussed next.

#### 2. NARMAX model

Although Volterra series can provide an intuitive representation for nonlinear systems, there are several critical limitations including i) it cannot represent severely nonlinear systems; ii) the order of the Volterra series expansion can be very high in order to achieve a good approximation accuracy; however iii) the estimation of high order Volterra kernel requires a large number of data and can be computationally very expensive. Nonlinear Autoregressive Moving Average Model with Exogenous Inputs (NARMAX) model [59, 61] has therefore been developed as an alternative to the Volterra series. NARMAX model normally contains a much smaller number of terms due to the inclusion of output delay terms, and its identification process is computationally more efficient. Similar to the Volterra series, a polynomial Nonlinear Autoregressive Model with Exogenous Inputs, NARX (the simplest NARMAX) model, can be expressed as a summation of a series of output terms with different orders of nonlinearity:

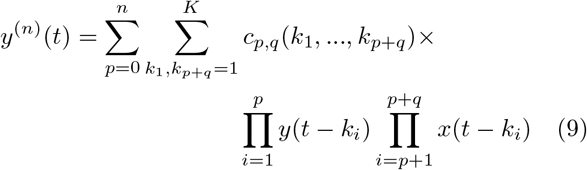

where *p* + *q* = *n, k_i_* = 1,…, *K*, and 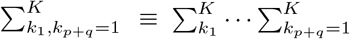. The number of model terms depends on the order of input and output (*q* and *p*) and the maximum lags (*K*). The NARX model structure and parameters are typically identified based on the forward regression with the orthogonal least squares (FROLS) method [62]. In cases where the system under study is stochastic with unknown coloured noise, noise moving average (MA) models should be employed to form a full NARMAX model. The identified model can be statistically validated using nonlinear correlation tests [63, 64].

A wide range of nonlinear systems can be represented by NARMAX method, including systems with exotic nonlinear behaviours such as subharmonics, bifurcations, and chaos, as observed in the human nervous system [65]. Until now, NARMAX methodology has been employed to develop dynamic models for nonlinear sensory processing circuit from spiking neuron data [66] as an improvement to the previous Volterra model-based studies [57], to investigate the somatosensory afferent pathways from muscles to the brain [67, 68]; as well as to study the corticothalamic nonlinear interactions during tremor active and resting states [9]. Apart from efficient time-domain predictive modelling, NARMAX also provides an essential base for the nonlinear frequency-domain analysis, nonlinear time-varying modelling, and nonlinear causality analysis to be discussed in the following sections.

#### 3. Dynamic causal modelling

Most of the effective connectivity models, e.g. linear and nonlinear autoregressive models, are directly identified from functional neurophysiological signals. However, sometimes it would be more accurate and meaningful to identify the causal interactions of the underlying neuronal activities at the level of neuronal dynamics [69]. The aim of dynamic causal modelling (DCM) [60, 70] is to infer such connectivity among brain regions (or sources) under different experimental factors or inputs. A DCM comprises typically two parts: a neuronal part that describes dynamics among neuronal sources and a measurement part that describes how the source dynamics generate measurable observations, e.g. EEG or MEG [71, 72]. Therefore, DCM can be naturally expressed as a nonlinear state-space model with hidden states denoting unobserved neuronal dynamics and the observation equation (e.g. the lead-field) assumed linearly in the states. The effective connectivity among those sources can be identified via Bayesian model selection and Bayesian inference of the neuronal model parameters. One strength of DCM is its biophysical and neuronal interpretation of how the neurophysiological signals are generated from the underlying neuronal system, through the generative or forward (state-space) modelling. Due to the complexity and computational cost of Bayesian model selection, DCM is more suitable to investigate the connectivity among pre-defined regions of interest, rather than exploratory analysis of relatively large brain or neural networks [73]. Compared to the hypothesis-driven DCM, the NARMAX or Volterra models are more flexible in terms of model structure identification and their direct frequency-domain mapping (to be discussed) is a powerful tool to study the nonlinear cross-frequency interactions between neurological regions.

#### 4. Other black-box neural nonlinear system identification methods

Apart from the aforementioned three important generic nonlinear model representations, other black-box modelling approaches have also been applied in the ‘neural system identification’ context. For example, artificial neural networks (ANNs), e.g. recurrent, multilayer perceptron, fuzzy, probabilistic neural networks, have often been used as an alternative to classical system identification models. ANNs have been applied to predict neural responses in visual cortex [74, 75], and to improve the prediction of synaptic motor neuron responses [76]. More recently, deep neural networks (DNNs), such as convolutional neural network (CNN) or recurrent neural network (RNN), are employed to model sensory neural responses, to understand neural computations and to learn feature spaces for neural system identification [77–81]. Nevertheless, in the current neuroscience literature, ANNs and DNNs are applied more towards automatic feature extraction and classification problems rather than traditional ‘system identification’. For instance, automatic detection and diagnosis of neurological disorders via a combination of ANN with other nonlinear feature extraction techniques such as approximate entropy and wavelet [82–85], or direct implementation of DNNs [86, 87]. Nonparametric Bayesian approaches like Gaussian process (GP) is closely related to ANN. GP has recently been used for system identification purpose [88–90] and applied to analyse neurophysiological signals [91], such as the use of GP modelling for EEG-based seizure detection and prediction [92] and heteroscedastic modelling of noisy high-dimensional MEG data [93]. Compared with ANN, GP can be applied to model datasets with small sample size and it has a relatively small number of hyperparameters. Additionally, due to its Bayesian nature, GP can incorporate prior knowledge and specifications into the modelling and can directly capture the model uncertainty. Another well-known system identification paradigm is the nonlinear state-space model [94, 95]. Its strength in dynamic (latent) state estimation and sequential inference process makes it a suitable candidate in the identification of certain neural systems. The state-space models have been applied to infer neural spiking activity induced by an implicit stimulus observed through point processes [96], to perform optimal decoding given multi-neuronal spike train data and tracking nonstationary neuron tuning properties (for a review, see [97]), and to perform source localization from neurophysiological signals like MEG and EEG [98, 99]. All of those black-box modelling approaches are usually flexible and accurate in quantifying complex and long-range nonlinear interactions. In comparison, the advantages of NARMAX and Volterra models are their modelling simplicity, interpretability of nonlinear interactions in the time-domain (e.g. the order of nonlinearity, phase delay), and frequency-domain mapping and analysis (e.g. energy transfer, intermodulations).

### B. Frequency-domain nonlinear system analysis: nonlinear frequency response functions

Many nonlinear effects, such as harmonics, intermodulations and energy transfer, can only be accurately and intuitively characterised in the frequency domain. Thus, it is important to ‘map’ the identified time-domain nonlinear models to the frequency domain for further analysis. A multidimensional Fourier transform of the *n*th-order Volterra kernel in (8) yields the so-called *n*th-order generalised frequency response function (GFRF), *n*th-*H_n_*(*f*_1_,⋯, *f_n_*), which is a natural extension of the linear frequency response function to the nonlinear case [59]. The output spectrum *Y*(*f*) of a nonlinear system can then be expressed as a function of the input spectrum *X*(*f*) and GFRF, known as the output frequency response function (OFRF) [59, 100]:

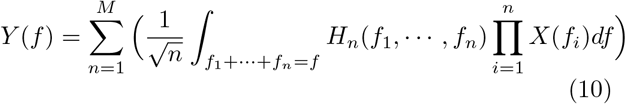

Compared with the Volterra series, the GFRFs can be more efficiently computed from the identified time-domain NARMAX model (9) and corresponding model parameters [101]. As shown in Figure 2, the peaks in 1st-order GFRF indicate the well-known ‘resonance frequencies’ of the linear part of the system; and the peaks (or ridges) in the 2nd-order GFRF would indicate nonlinear harmonics (*f*_1_ + *f*_2_ when *f*_1_ = *f*_2_) or inter-modulation effects (*f*_1_ ± *f*_2_ when *f*_1_ ≠ *f*_2_) in the output spectrum, and so on. Since 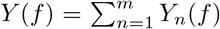, the *n*th OFRF *Y_n_*(*f*) represents the *n*th-order (linear or nonlinear) contribution from the input to the output spectrum. Practically, by comparing the OFRF with the spectrum of the output signal, obtained from a classical nonparametric estimation such as fast Fourier transform, one can also ‘validate’ the accuracy of the time-domain modelling in addition to the aforementioned NARMAX model validation [9].

**FIG. 2:**
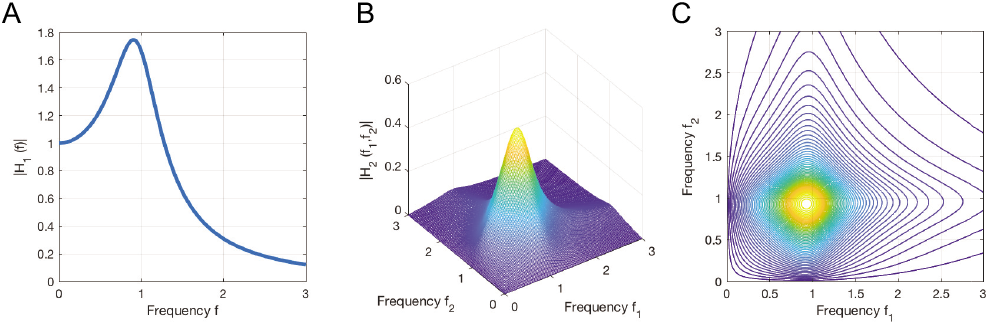
The GFRFs of an exemplar nonlinear system. (A) The linear 1st-order GFRF, *H*_1_(*f*), shows a ‘resonance’ peak at *f* = 0.9Hz; (B) and (C) the 3-D and contour plots of the 2nd-order GFRF, *H*_2_(*f*_1_, *f*_2_). It shows a peak at *f*_1_ = 0.9*Hz*, *f*_2_ = 0.9*Hz*, which indicates harmonics at 2*f* = *f*_1_ + *f*_2_ = 1.8*Hz* can be introduced in the output spectrum if input contains a 0.9Hz component.

NARMAX-based frequency-domain analysis method has been applied to quantify the dynamic characteristics of nonlinear sensory processing circuit models from spiking neuron data [66], the cross-frequency interactions in the corticothalamic loops with respect to tremor [9], and the characterisation of epileptic brain states [64]. More details will be discussed in the Section VI.

### C. Time-varying nonlinear system identification

Many neurological subsystems are inherently nonstationary, since the brain is a dissipative and adaptive dynamical system [102, 103]. Modelling and identification approaches of nonstationary processes have been well developed for linear systems, i.e. linear time-varying (LTV) systems. They are primarily based on adaptive recursive methods, such as recursive least squares, least mean squares, and the Kalman filter [104], or based on a finite basis function expansion of the time-varying coefficients [105–108]. The identification of a nonlinear time-varying system is more sophisticated. The primary difficulty is how to effectively and simultaneously resolve the nonlinear model structure selection and the time-varying coefficient estimation. Approaches based on time-varying Volterra series combining artificial neural networks [109], principal dynamic modes [110], or sliding-window strategy [111], have been proposed. However, the model structure selection is still an unsolved issue, and the identification and frequency-domain analysis are computationally costly.

A better strategy is to extend the basis function expansion approach, originally proposed for LTV identification, to nonlinear time-varying cases [112]. The timevarying (TV) parameters in TV-NARX models are first expanded using multi-wavelet basis functions, and TV nonlinear model is transformed into an expanded time-invariant model; the challenging TV model selection and parameter estimation problem can then be solved using the computational efficient FROLS algorithm. To accommodate the stochastic perturbations or additive coloured noise, this procedure can also be extended to more general TV-NARMAX models by introducing a modified extended least squares (ELS) algorithm [113]. Several modifications to the TV-NARX model has recently been proposed using different basis functions or model selection procedure [114, 115]. The corresponding frequency-domain analysis for non-linear time-varying systems has also been developed based on the identified time-domain TV-NARX or TV-NARMAX model and the TV-GFRF concepts [113, 116]. By fitting TV-NARX models to two fragments of intracranial EEG recordings measured from epileptic patients, the corresponding frequency-domain analysis (i.e. TV-GFRF and TV-OFRF) shows the nonlinear energy transfer effect – the underlying neural system transfers the energy from lower frequencies to higher frequencies as the seizure spreading from the left to the right brain regions over time [116, 117].

An overview of the NARMAX model-based system identification framework, including both time-invariant (NARMAX) and time-varying (TV-NARX) modelling along with corresponding frequency-domain analysis to neurophysiological signal analysis, is summarised in Figure 3.

**FIG. 3:**
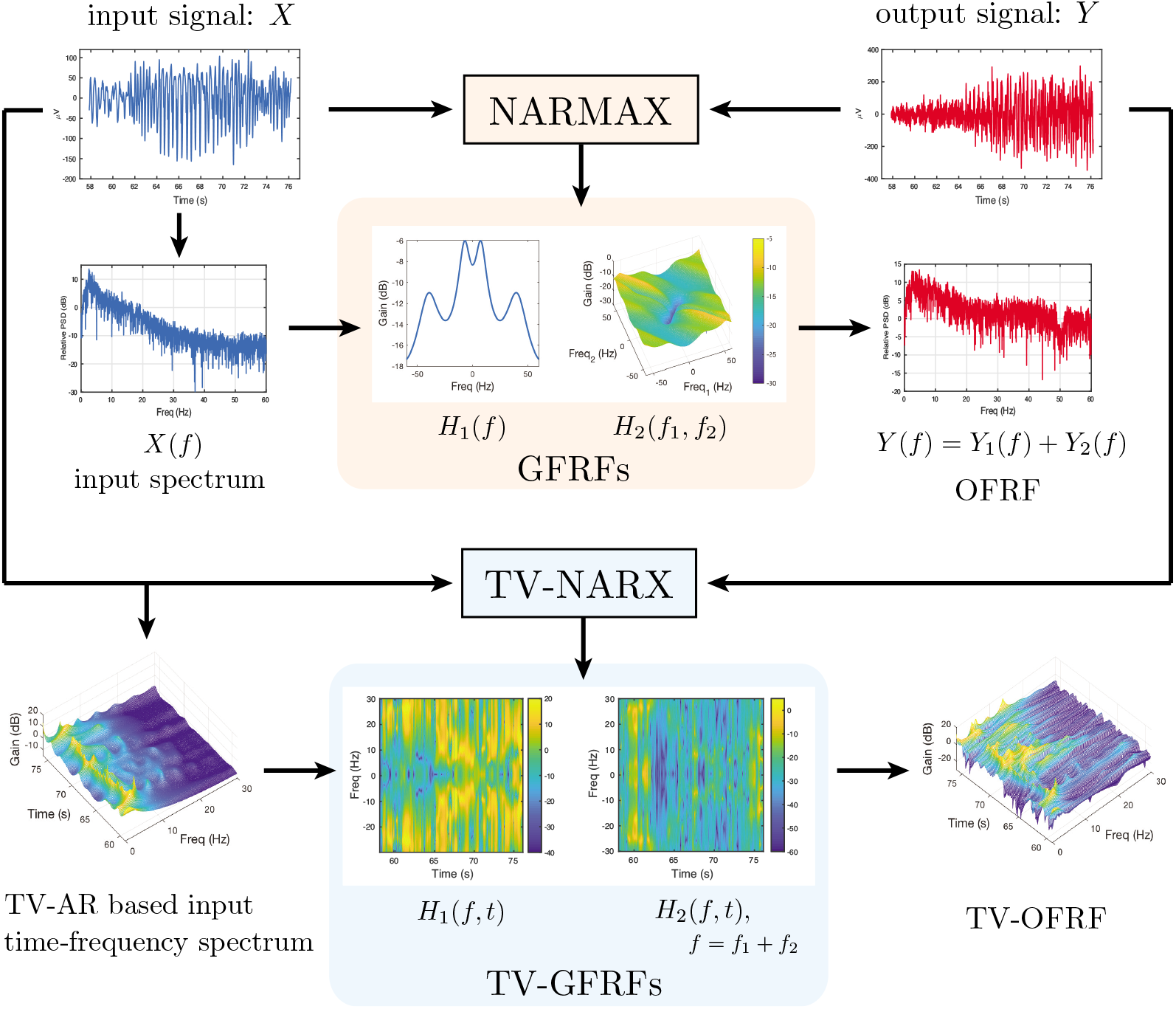
Analysing neurophysiological signals uses nonlinear time-invariant and time-varying system identification and corresponding frequency-domain analysis methods. The upper part of the diagram illustrates the nonlinear time-invariant modelling: first a NARMAX model is identified from the input and output neurophysiological signals (e.g. EEG, EMG, MEG, LFP); this time-domain model is then mapped to the frequency-domain with GFRFs (i.e. *H*_1_(*f*), *H*_2_(*f*),…), and the OFRF (*Y*(*f*) = *Y*_1_(*f*) + *Y*_2_(*f*),…) can be computed from the input spectrum and GFRFs. The lower part of the diagram shows the nonlinear time-varying system identification using a TV-NARX model, and the identified time-varying model can then be mapped to the time-frequency domain with (averaged) TV-GFRFs (i.e. *H*_1_(*f, t*), *H*_2_(*f, t*),…). The TV-OFRF can therefore be computed from a combination of the input time-frequency spectrum and the TV-GFRFs.

## V. NONLINEAR NEURAL CONNECTIVITY ANALYSIS

The communication between different neuronal populations which have very different firing behaviours can result in nonlinear neural connectivity, showing neural coupling across two or more different frequencies. To quantitatively study such a ‘cross-frequency coupling’, this section reviews recent advances in nonlinear neural functional and effective connectivity analysis.

### A. High-order spectrum and nonlinear coherence

The power spectra and coherence discussed in Section III A are Fourier transforms of the auto- and cross-correlations of signals, hence they are only linear frequency-domain measures. Practically these measures cannot detect certain nonlinear effects such as quadratic moments in or between signals that have a zero mean [59]. Higher-order spectral analysis has been developed to detect nonlinear coupling between spectral components [118]. For example, the most widely used bispectral analysis is the frequency-domain mapping of the third-order statistics. It can be used to quantify the quadratic nonlinear interactions, i.e. quadratic phase coupling. The bispectrum or bicoherence (and the bivariate cross-bispectrum or cross-bicoherence) analysis is well-known in engineering signal processing, whereas it has only relatively recently been applied to study the nonlinear interactions in neurophysiological signals [119–121]. For example, bispectral measures were used to detect long-range nonlinear coupling and synchronization in healthy subjects from human EEG [120, 122], to characterise and predict epileptic seizures [123], and to study the nonlinear interactions between different frequency components related to Parkinson’s disease and tremor [9, 124, 125].

However, bispectrum or bicoherence cannot detect nonlinearity beyond second order, such as the higher-order harmonics and intermodulation effects, or the subharmonic coupling. A generalised nonlinear coherence analysis framework has therefore been proposed, based on two different nonlinear mappings from the input to the output of an ‘open-loop’ system in the frequency domain [6]:

1. n:m Mapping: to measure harmonic or subharmonic coupling related to individual input frequencies. 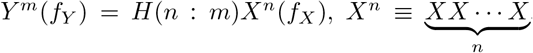, where the output frequencies (*f_Y_*) are related to the input frequencies (*f_X_*) by the ratio *n/m* (*n* and *m* are co-prime positive integers), and *H*(*n*: *m*) is the n:m mapping function. The n:m mapping can generate cross-frequency (e.g. harmonic *m* = 1 or subharmonic *m* > 1) coupling between the input and the output.
2. Integer Multiplication Mapping: to quantify inter-modulation coupling among multiple (≥ 2) input frequencies.

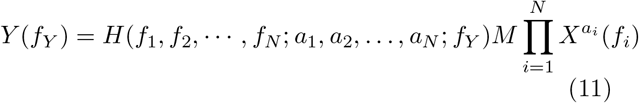

where *f_Y_* = *a*_1_ *f*_1_ + *a*_2_ *f*_2_ + ⋯ + *a_N_ f_N_*. The *M* is the corresponding multinomial coefficient, and *H*(*f*_1_, *f*_2_, ⋯, *f_N_*; *a*_1_, *a*_2_,…, *a_N_*; *f_Y_*) indicates amplitude scaling and phase shift from the input to the output. According to these two different types of nonlinear mapping, Yang and colleagues proposed two basic metrics for quantifying nonlinear coherence: (i) n:m coherence and (ii) multi-spectral coherence [6].

#### 1. n:m coherence

The n:m coherence is a generalized coherence measure for quantifying nonlinear coherence between two frequency components of the input *X*(*f*) and the output *Y*(*f*) [6]:

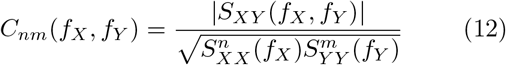

where *n*: *m* = *f_Y_*: *f_X_*. *S_XY_*(*f_X_, f_Y_*) =< *X^n^*(*f_X_*)(*Y^m^*(*f_Y_*))* > is the n:m cross-spectrum, with <·> represents the averaging over repetitions. 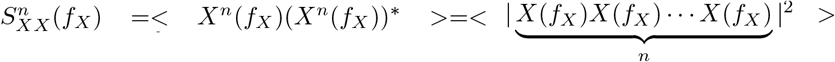 is the *n*th-order auto-spectra. According to Cauchy-Schwarz-inequality, we have:

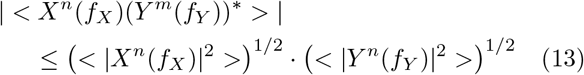

Thus, n:m coherence is bounded by 0 and 1, where one indicates that two signals are perfectly coupled for the given frequency pair (*f_X_, f_Y_*).

A simplified version of n:m coherence that considers only the phase relation between the input and the output is known as n:m phase synchronization index [126]. The n:m coherence and n:m phase synchronization index has been widely applied to neuroscience research to investigate nonlinear functional connectivity in different brain regions [127, 128], as well as the nonlinear connectivity between the brain and muscles [36].

#### 2. Multi-spectral coherence

Multi-spectral coherence measures the multi-frequency nonlinear coupling generated by the integer multiplication mapping [6]. It is defined as:

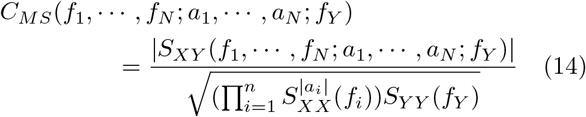

where 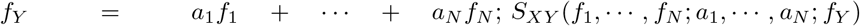 is the high-order cross-spectrum between *X* and *Y*, and equal to 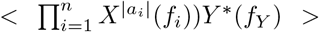. Here ‘*’ indicates the complex conjugate. When there are only two input frequencies, the multi-spectral coherence is degraded to the bicoherence [129]. The multi-spectral coherence or bicoherence has been applied to study the nonlinear behaviours in visual [130], auditory [131] and somatosensory systems [6], which are thought to be associated with neural coding and functional integration of various sensory inputs [132].

A simplified version of multi-spectral coherence that considers only the phase relation between the input and the output is known as multi-spectral phase coherence [133]. Similarly, there is a degraded measure, named biphase locking value [134], for the case involving only two input frequencies. The advantage of multi-spectral phase coherence or bi-phase locking value is that it allows precise estimation of time delay in the nervous system based on the relative phase relationship between the input and output [133, 135]. The multi-spectral phase coherence or bi-phase locking value has been previously used to determine neural transmission delays in the human visual system [130] and the stretch reflex loops [36].

### B. Nonlinear causality analysis: system identification based approaches

#### 1. Time-domain analysis

In terms of effective connectivity, classical linear Granger causality analysis (GCA) (as discussed in Section III B) may provide misleading results when used to analyse EEG/MEG or EMG signals, as the possible nonlinear interactions within a neural system are not modelled explicitly by simply using linear regression models. The Granger causality definition has been extended to nonlinear cases in the time domain, based on nonparametric methods [136, 137], radial basis functions [138], kernel methods [139], local linear models [140]. DCM [60] (see Section IV A 3) was developed to accommodate both linear and nonlinear causal effects using a dynamic state-space model, and the effective connectivity among hidden states (unobserved neuronal dynamics) can be identified via Bayesian inference. Information-theoretical effective connectivity measures have also been proposed, such as the bivariate transfer entropy (TE) [141, 142] and phase transfer entropy (PTE). TE is a model-free measure, which compares two conditional probabilities using the Kullback-Leibler divergence - the amount of uncertainty in the future of target signal *Y* conditioned only on the target’s past and the future of *Y* conditioned on the past of both its own *Y* and the source *X*, in a conceptually similar way as the GCA. A more recent work [143] generalised the TE method by using multivariate transfer entropy, which can overcome the problems of inferring spurious or redundant causality and missing synergistic interactions between multiple sources and target.

Another strategy to implement nonlinear granger causality under a system identification framework is to use NARX models [144, 145], by calculating the relative predictability improvement obtained from the NARX model compared to the corresponding NAR model. More importantly, compared to other nonlinear causality measures (e.g. nonparametric or information-theoretic measures), the advantage of using NARX-based causal inference [145] is that one can easily separate the linear and nonlinear causal influence, for example from an input *X* to an output *Y*. After fitting a polynomial NARX model with the form (9), the linear causality can still be calculated from (4) based on the linear part of the NARX model, while the nonlinear causal influence from *X* to *Y* can be defined as:

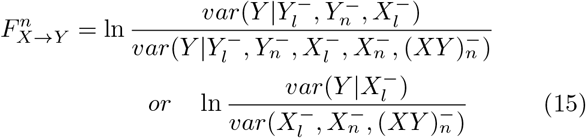

Here 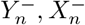 and 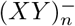 denote the sets of all nonlinear delayed terms of *Y, X* and nonlinear product terms *XY*. This nonlinear causality measure can also be generalised to nonlinear time-varying systems, by computing similar linear and nonlinear causality indices based on the identified TV-NARX models (as described in Section IVC), as proposed in [145, 146].

#### 2. Frequency-domain analysis

In the frequency domain, linear Granger causality measures, such as PDC, DTF and spectral Granger causality, can all be expressed as a function of the elements in the coefficient matrix or its inverse the transfer function matrix of the corresponding linear ARX models (3). By identifying the link between the PDC and the FRFs of the corresponding linear ARX models (as described in the Section III B), a new nonlinear PDC (NPDC) measure has been proposed [64] by generalising the spectrum decomposition with respect to a nonlinear NARX model in a similar way as the linear case. The NPDC from *X* to *Y* can then be expressed as a direct generalization of the linear PDC:

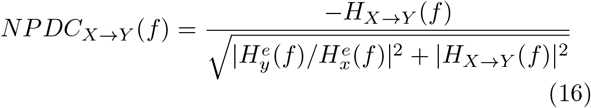

Here, the *H*_*X*→*Y*_(*f*) is the ‘nonlinear FRF’ which replaces the FRF in the linear PDC (7), and 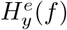 and 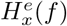 are the error-driven GFRFs correspond to the restricted NAR models with respect to *Y* and *X* as discussed in [64]. The NPDC measures both linear and nonlinear causal influences from *X* to *Y*. The linear causal effects can be interpreted as a special case of (16) by only considering the 1st-order nonlinear FRFs of NARX (i.e. *H*_1, *X*→*Y*_(*f*)) and NAR (i.e. 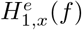 and 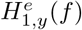) models.

This new NPDC measure has recently been applied to predict epileptic seizures from EEG data [147] by advancing the construction of functional brain networks, nonlinear feature selection and classification. This new nonlinear causality measure helps to provide better prediction accuracy compared to other standard graph theory or nonlinear classification based methods. A nonlinear generalization of Geweke’s spectral Granger causality has also been proposed [117] using the NARX methodology.

## VI. NEUROLOGICAL AND CLINICAL APPLICATIONS

Movement, sensation and cognition arise from the cumulative activity of neurons within neural circuits and across distant, macroscale networks in the nervous system. Although the behaviour of an individual neuron has been investigated and well understood for decades, the mechanisms underlying neural communications between macroscale neural networks are still yet to better understand. Newly developed nonlinear system identification approaches allow us to investigate neural communications from large-scale neural activities measured by EEG, MEG and EMG, with the most recent application examples discussed below.

### A. Nonlinear cortical response to somatosensory inputs

The human somatosensory system is highly nonlinear [11]. Previous studies applied periodic sinusoid tactile stimulations to the index finger and measured the cortical response, where they found harmonic and subharmonic patterns in the response [5, 148]. Several recent studies used sum-of-sinusoid stimulations to the wrist joint and found not only harmonics and subharmonics but also intermodulation patterns [6, 29]. The majority of intermodulation responses presented the second-order nonlinearity, which is the sum or the difference between input frequencies [8]. These findings indicate that the nonlinearities in the somatosensory system allow the functional integration of input signals at different frequencies, and they can be transmitted in different somatosensory ascending pathways.

Yang and colleagues recently built a hierarchical neural network based on known neuroanatomical connections and corresponding transmission delays in neural pathways to model the cortical response to somatosensory input [67]. The proposed computational model contains a neural layer at the thalamus that integrates the inputs from different ascending pathways, including Ia and II afferents. The computational model well captured the majority of the cortical response to the given somatosensory inputs, indicating the functional integration of different somatosensory input signals may occur at the thalamus and is then transmitted to the cortex via the thalamocortical radiation.

### B. Tremor: nonlinearity in the thalamocortical loop

Essential tremor is a common neurological movement disorder widely considered to have a centrally-driven origin. There is both neurophysiological and clinical evidence of thalamic involvement in the central oscillatory network generating essential tremor [149–151]. Local field potential (LFP) recordings of thalamic ventralis intermedius (Vim) nucleus show a strong linear correlation with the contralateral EMG during tremor [150]. Some studies using EEG and MEG suggest that the sensorimotor cortex is also part of the central tremor-related oscillatory network, with significant coupling in some cases with the contralateral tremorgenic EMG [152–154]. Despite a well-established reciprocal anatomical connection between the thalamus and cortex, the functional association between the two structures during ‘tremor-on’ periods had not been extensively defined.

He and co-authors [9] investigated the functional connectivity among cortical EEG, thalamic (Vim) LFPs and contralateral EMG signals over both ‘tremor-on’ and ‘tremor-off’ states, using linear coherence and nonlinear bispectral analysis methods. In addition to expected strong coherence between EMG and thalamic LFP, nonlinear interactions (i.e. quadratic phase coupling) at different frequencies, i.e. low frequency during tremor on and higher frequency during tremor off, in LFPs have been reported. More importantly, by using the NARX-based nonlinear system identification and frequency-domain analysis (as described in Section ‘IV B’), two distinct and non-overlapping frequency ‘channels’ of communication between thalamic Vim and the ipsilateral motor cortex were identified, which robustly defined the ‘tremor-on’ versus ‘tremor-off’ states. Longer corticothalamic nonlinear phase lags in the tremor active state were also uncovered, suggesting the possible involvement of an indirect multi-synaptic loop. This work demonstrates, for the first time, the importance of cross-frequency nonlinear interactions between the cortex and the thalamus in characterising the essential tremor.

### C. Nonlinear analysis for determining motor impairment in stroke

After a stroke, damage to the brain increases the reliance on indirect motor pathways resulting in motor impairments and changes in neural connectivity between the brain and muscles. A hallmark of impairments post-stroke is a loss of independent joint control that leads to abnormal co-activation between shoulder, arm and hand muscles, known as the upper limb synergy [155]. The upper limb synergy is thought to be caused by progressive recruitment of indirect motor pathways via the brainstem following a stroke-induced loss of corticospinal projections [156]. Thus, a neural connectivity measure that quantifies the recruitment of these indirect motor pathways would be crucial to evaluate post-stroke motor impairments. Recent model-based simulation and clinical studies indicate that the increased usage of indirect motor pathways enhances nonlinear distortion of motor command transmissions, which leads to stronger nonlinear interaction between the brain and muscles [38, 40]. The ratio of nonlinear interaction over linear interaction, known as the nonlinear-over-linear index (N-L Index), has been reported to be associated with the relative ratio of the recruitment of indirect versus direct motor pathways [40]. This new measure may facilitate the future determination of the effect of new therapeutic interventions that aim to optimise the usage of motor pathways, and thus facilitate the stroke recovery.

### D. Epilepsy

It has been widely recognised that epileptic seizures are highly nonlinear phenomena, due to low dimensional chaos during epileptic seizure or transitions between ordered and disordered stats [157]. Currently, the treatment mainly relies on long-term medication with antiepileptic drugs or neurosurgery, which can cause cognitive or other neurological deficits. New treatment strategies such as on-demand therapies during the preseizure (preictal) state or electrical stimulation are therefore needed. A vital part of this new on-demand strategy is the accurate and timely detection of the preictal state, even seconds before seizure onset [158]. A range of univariate, bivariate and multivariate linear and nonlinear measures have been developed for the characterisation and detection or prediction of epileptic brain states and achieving a better understanding of the spatial and temporal dynamics of the epileptic process [158, 159]. There is a comprehensive review of using different parametric and nonparametric nonlinear features (in time, frequency and time-frequency domains) for the automated epilepsy stage detection and classification [160].

Given the current challenges in epilepsy detection and diagnostics [158, 161], e.g. to improve the understanding of brain dynamics and mechanisms during the seizure-free interval and seizure initiation and termination, there is a great need to develop more accurate nonlinear methods to improve the detectability of directional interactions in underlying functional and anatomical networks. Developing new nonlinear system identification and nonlinear causality measures are therefore crucial. A nonlinear causality measure, partial transfer entropy [162], has been applied to analyse the EEG of epileptic patients during preictal, ictal and postictal states. It can provide better detection of causality strength variations compared to linear PDC. An adaptive nonlinear Granger causality measure was also proposed [163] and applied to LFP data (intracranial EEG in cortex and thalamus) in rats. It was reported to provide more sensitive detection of changes in the dynamics of network properties compared to linear Granger causality. The recently proposed nonlinear frequency-domain causality measure NPDC [64] (as reviewed in Section ‘VB2’) has been applied to analyse EEG recordings of two bipolar channels from a patient with childhood absence epilepsy. It shows this nonlinear measure can detect extra frequency-domain causal interactions compared to standard linear PDC.

## VII. CONCLUSIONS AND PERSPECTIVES

The complexity and nonlinearity of neural systems require advanced system identification techniques to understand their properties and mechanisms better. This review investigated the links between connectivity analysis and system identification, as well as recent progress of nonlinear system identification of neural systems. With the state-of-the-art examples of clinical applications, we argued that nonlinear dynamic modelling and corresponding connectivity analysis allows new insights into the underlying neural functions and neuropathological mechanisms of the abnormality caused by various neurological disorders. These novel approaches may well facilitate the development of new precision diagnostic tools and brain-computer interface (BCI) techniques [164–166], and therefore improve the diagnosis and treatment of neurological disorders as well as restore communication and movement for people with motor disabilities.

Compared to the linear system identification and iso-frequency connectivity analysis, nonlinear dynamic modelling and cross-frequency analysis are much more complicated. Such complexity brings challenges but also research opportunities. Potential future work includes: i) further developing multivariate system identification techniques and corresponding multivariate nonlinear frequency-domain analysis and causality analysis measures. Most existing nonlinear system identification based (time or frequency-domain) analysis or causality analysis are primarily bivariate, which limits nonlinear analysis to the only pre-defined local brain or neural regions. New multivariate system identification (e.g. multivariate nonlinear regression modelling) or inference approaches would generalise the existing nonlinear connectivity analysis to larger neural networks, although developing efficient model selection and reducing the computational cost would be challenging tasks; ii) many neuronal systems or interactions are in nature nonstationary and nonlinear, how to automatically distinguish the nonlinearity and time-varying effects (nonstationarity) via novel system identification technique is still an open and important research topic, although significant progress has been made so far (as reviewed in Section IVC); iii) machine learning and deep learning techniques have recently been applied to Granger causality analysis [167–169], an interesting future work is to further explore and combine the advantages of deep learning, e.g. accurate quantification of complex and long-range nonlinear interactions, and nonlinear system identification approaches to study the nonlinear causal interactions in complex neural networks; iv) using nonlinear system identification techniques to extract new nonlinear features for the BCI; v) apart from those clinical applications described in Section VI, the importance of nonlinearity in understanding and characterising other important neurological disorders, e.g. Parkinson’s disease and Alzheimer’s disease [170–173], has been reported recently. Therefore nonlinear system identification approaches will have great potential in developing new diagnostic tools for those primary neurological disorders that affect a large population worldwide.

